# Oligomer-to-Monomer Transition Underlies the Chaperone Function of AAGAB in AP1/AP2 Assembly

**DOI:** 10.1101/2022.03.11.483825

**Authors:** Yuan Tian, Ishara Datta, Rui Yang, Chun Wan, Bing Wang, Huan He, Suzhao Li, Jingshi Shen, Qian Yin

**Author notes:** These authors contributed equally to the work (Yuan Tian, Ishara Datta). Correspondence (J.S.); (Q.Y.).

## Abstract

Assembly of protein complexes is facilitated by assembly chaperones. Alpha and gamma adaptin binding protein (AAGAB) is a chaperone governing the assembly of the heterotetrameric adaptor complexes 1 and 2 (AP1 and AP2) involved in clathrin-mediated membrane trafficking. Here, we found that before AP1/2 binding, AAGAB exists as a homotetramer. AAGAB tetramerization is mediated by its C-terminal domain, which is critical for AAGAB stability and is missing in mutant proteins found in patients with the skin disease punctate palmoplantar keratoderma type 1 (PPKP1). We solved the crystal structure of the tetramerization domain (TD), revealing a dimer of dimer assembly. Interestingly, AAGAB uses the same TD to recognize and stabilize the γ subunit in the AP1 complex and the α subunit in the AP2 complex, forming binary complexes containing only one copy of AAGAB. These findings demonstrate a dual role of TD in stabilizing resting AAGAB and binding to substrates, providing a molecular explanation for disease-causing AAGAB mutations. The oligomerization state transition mechanism may also underlie the functions of other assembly chaperones.

## Introduction

Formation of protein complexes is facilitated by assembly chaperones that transiently interact with individual subunits and assembly intermediates to prevent aggregation and ensure correct assembly (Ellis, 2013). Substrate-specific assembly chaperones have been identified for a number of protein complexes including ribosomes, nucleosomes, proteosomes, and spliceosomes (Ellis, 2013).

In clathrin-mediated membrane trafficking, alpha and gamma adaptin binding protein (AAGAB, also known as p34) is an assembly chaperone required for the formation of the clathrin adaptors AP1 and AP2 (Gulbranson et al., 2019; Wan et al., 2021). Clathrin coats drive the budding of vesicles from the endosome, the plasma membrane, and the *trans*-Golgi network (Kirchhausen et al., 2014; Mettlen et al., 2018; Traub and Bonifacino, 2013). Clathrin relies on adaptors to recruit cargo proteins to vesicle budding sites (Kaksonen and Roux, 2018; Page et al., 1999). Two predominant clathrin cargo adaptors are the AP1 complex involved in trafficking from the endosome and the *trans*-Golgi network, and the AP2 complex regulating clathrin-mediated endocytosis (CME) (Traub, 1997; Traub and Bonifacino, 2013). Both AP1 and AP2 are heterotetrameric complexes with two large subunits (γ/α and β), one medium subunit (μ), and one small subunit (σ) (Collins et al., 2002; Conner and Schmid, 2003; Heldwein et al., 2004; Hollopeter et al., 2014; Pearse and Robinson, 1984; Ren et al., 2013).

In AAGAB-assisted AP1 assembly, AAGAB first binds to the γ subunit (gamma adaptin) to form an AAGAB:γ binary complex, which then recruits the σ subunit to form an AAGAB:γ:σ ternary complex. In these complexes, AAGAB stabilizes the γ and σ subunits as well as the γ:σ hemicomplex. Subsequently, β and μ subunits displace AAGAB, leading to the formation of the AP1 complex (Gulbranson *et al*., 2019; Wan *et al*., 2021). AAGAB regulates the assembly of the AP2 adaptor through a similar mechanism (Gulbranson *et al*., 2019). Without the assistance of the assembly chaperone AAGAB, AP1 and AP2 complexes fail to form, resulting in degradation of their subunits and membrane trafficking defects (Gulbranson *et al*., 2019). Autosomal dominant mutations in the *AAGAB* gene cause punctate palmoplantar keratoderma type 1 (PPKP1, also known as Buschke-Fischer-Brauer disease), a skin disease characterized by lesions on palms and soles (Giehl et al., 2012; Kono et al., 2017; Nomura et al., 2015; Pohler et al., 2012; Pohler et al., 2013).

It remains unclear how AAGAB recognizes AP1/2 subunits and how AAGAB-AP1/2 interactions promote AP1/2 adaptor assembly. In this work, we uncovered that AAGAB itself tetramerizes in the absence of AP1/2, mediated by a highly conserved C-terminal domain we henceforth designated tetramerization domain (TD). Without TD, AAGAB becomes unstable and is degraded in the cell. We solved the crystal structure of the TD tetramer, which revealed a tetramerization interface formed by dimerization of two extended anti-parallel alpha helical dimers. Interestingly, AAGAB uses this exact TD to bind and stabilize AP1γ and AP2α subunits. There is just one copy of AAGAB in the AAGAB:γ and AAGAB:α binary complexes, suggesting that AAGAB tetramers dissociate into monomers upon AP1/2 binding. Structure-guided mutations in the TD region disrupt both AAGAB tetramerization and its binding to the γ subunit of AP1 and the α subunit of AP2. Consistent with these *in vitro* observations, the TD mutations disrupt the assembly of AP1 and AP2 adaptors and impair clathrin-mediated trafficking in the cell. Since *AAGAB* mutations in PPKP1 patients usually lead to truncated mutants lacking the TD, our findings demonstrate that the mutants are unable to engage in AP1γ and AP2α interactions. Taken together, our study unveiled that the AP1/2-binding region of AAGAB oligomerizes prior to AP1/2 association to stabilize the substrate-free state. We suggest that the oligomer-to-monomer transition mechanism also operates in other chaperone-assisted assembly processes.

## Results

### AAGAB exists as a tetramer in its resting state

AAGAB is a cytosolic protein with two conserved regions – an N-terminal G protein-like domain (GD, residues 1-177) and a C-terminal domain (CTD, residues 258-315) without any known homologous structure (Fig. 1a). The two conserved domains are connected by a ~70-residue non-conserved linker (Fig. 1a). To investigate its biochemical and structural characteristics, we expressed and purified full-length (FL) AAGAB protein from *E. coli*. The purified protein exhibited excellent purity and homogeneity based on the size exclusion chromatography (SEC) profile and Coomassie blue-stained SDS-PAGE gel (Figs. 1a, 1b). Interestingly, we observed that FL AAGAB eluted at a position much earlier than its calculated molecular weight (MW) of 34.6 kDa. The elution position at 13.4 mL corresponded to an experimental MW of 144.2 kDa, which suggests a tetrameric assembly (with a calculated MW of 138.4 kDa) (Fig. 1a). Serial dilution of AAGAB at concentrations ranging from 1.26 to 0.14 mg/mL did not change AAGAB elution position from the Superdex 200 column (Fig. 1a), indicating that the tetrameric assembly is stable even at the lowest concentration examined.

**Fig. 1.**
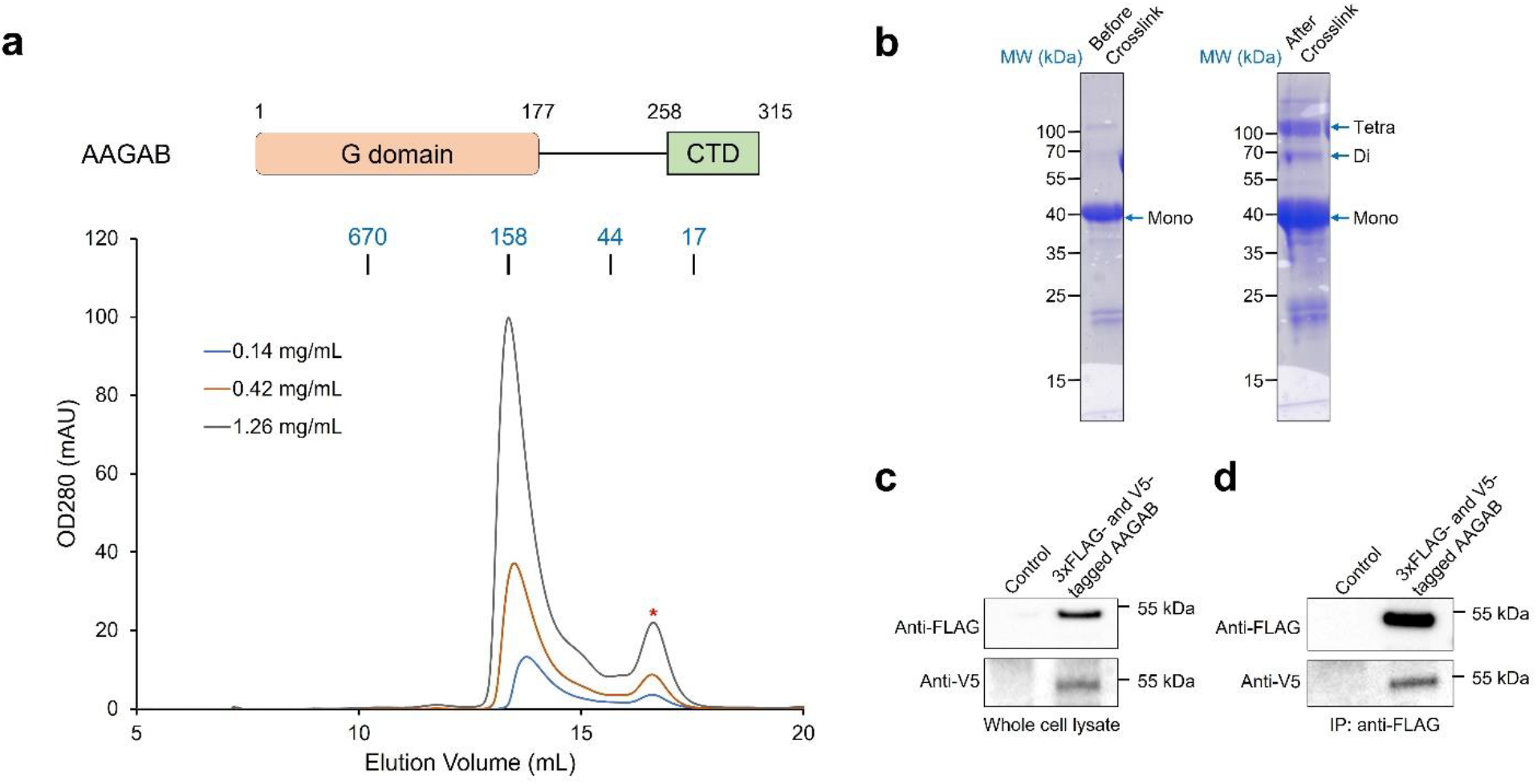
AAGAB exists as a tetramer in its resting state. **a** Domain architecture of FL AAGAB protein (top panel) and size-exclusion chromatography profiles of AAGAB protein at three 3-fold serial dilutions from a Supderdex200 increase 10/300 column. Starting and ending residue numbers and elution positions of protein standards with known MW are labeled above. The small peak marked by the red asterisk is from a contaminating protein. **b** Coomassie blue-stained gels showing AAGAB proteins before and after crosslinking. A new protein band consistent with tetramer size appears after crosslinking, in addition to the non-crosslinked monomer band. A minor new band is likely from the crosslinked AAGAB dimer. **c** Representative immunoblots showing the total expression of 3xFLAG- and V5-tagged AAGAB in *AAGAB* KO HeLa cells. Control: cells transfected with an empty vector. **d** Representative immunoblots showing the interaction between 3xFLAG- and V5-tagged AAGAB in the cells. The 3xFLAG-AAGAB protein was immunoprecipitated using anti-FLAG antibodies out of the whole cell lysates from **c** and the presence of 3xFLAG - AAGAB and V5-AAGAB in the immunoprecipitated products were measured by immunoblotting.

In parallel, we treated FL AAGAB with a lysine-specific crosslinking reagent and examined the crosslinked proteins using SDS-PAGE. As expected, denatured non-crosslinked FL AAGAB migrated as monomers on SDS-PAGE (Fig. 1b). Crosslinked AAGAB, however, exhibited a major new band consistent with a tetrameric MW (Fig. 1b). These crosslinking data are consistent with the results of the SEC experiments. We therefore conclude that AAGAB exists as a tetramer prior to AP1/2 binding.

Next, we examined whether AAGAB oligomerizes in the cell. AAGAB was N-terminally tagged with either a V5 or 3xFLAG epitope, which does not interfere with AAGAB function in the cell (Gulbranson *et al*., 2019). V5- and 3xFLAG-tagged AAGAB proteins were co-expressed in *AAGAB* knockout (KO) HeLa cells (Fig. 1c). We then performed co-immunoprecipitation (co-IP) using anti-FLAG antibodies. We observed that V5-tagged AAGAB was co-precipitated with 3xFLAG-tagged AAGAB (Fig. 1d), suggesting that AAGAB oligomerizes in the cell. These data are consistent with the *in vitro* results and further support the notion that AAGAB forms a tetramer in its resting state.

### AAGAB tetramerization is mediated by its CTD

To gain insights into the tetramerization interface(s), we performed mass spectrometry to analyze the lysine-specific crosslinked FL AAGAB proteins. We identified multiple intermolecular crosslinks between AAGAB chains. Interestingly, all crosslinked lysines mapped exactly to the short conserved CTD of AAGAB (Fig. 2a, 2b, Supplementary Table 1). All seven lysines in the CTD were crosslinked to other lysines from another AAGAB chain, whereas none of the seven lysines in the GD was crosslinked (Fig. 2a, 2b, Supplementary Table 1). These crosslinking results suggest that the CTD mediates the tetramerization of AAGAB.

**Fig. 2.**
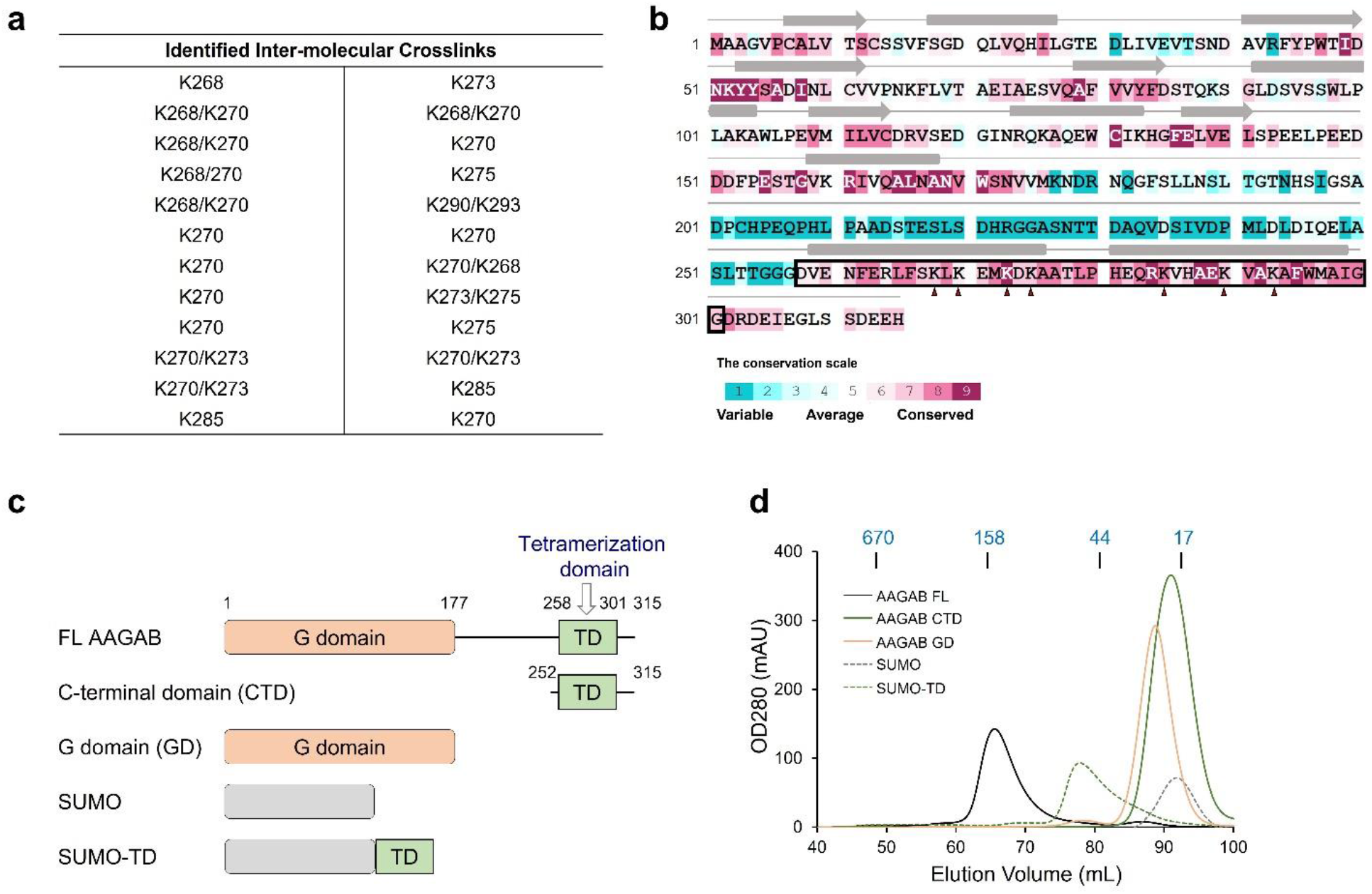
AAGAB tetramerization is mediated by its C-terminal tetramerization domain (TD). **a** Crosslinking mass spectrometry (XL-MS) mapped AAGAB interfacial residues solely to the TD. **b** Sequence analysis of the human AAGAB protein. The amino acids are colored based on their conservation levels. Predicted secondary structural elements are marked on top of the sequence: cylinder: α helix; arrow: β strand. The tetramerization domain (TD, residues 258-301) is highlighted in the black outline box. Crosslinked lysines identified in **a** are marked by red arrowheads. **c** Diagrams of FL, CTD (residues 252-315), GD (residues 1-177), and the SUMO-tagged TD (residues 258-301) of human AAGAB. **d** SEC profiles of the proteins depicted in **c** from a HiLoad 16/600 Superdex 200 column. Elution positions of protein standards with known MW are marked on the top. GD is monomeric (calculated MW of 19.7 kDa vs experimental MW of 23.3 kDa).

Both the conserved GD and CTD of AAGAB are predicted to fold into ordered structures, whereas the non-conserved linker region in the middle is predicted to be unstructured (Fig. 1a, 2b). We expressed the two structured domains individually in *E. coli* and purified them in a similar manner as FL AAGAB (Fig. 2c). The GD was monomeric based on its elution position on SEC (Fig. 2d). Interestingly, the CTD of AAGAB (residues 252-315) exhibited an estimated MW of 19.4 kDa on SEC (Fig. 2d), consistent with a tetrameric state. Based on our secondary structure prediction, the CTD contains a long alpha helix flanked by flexible loops (Fig. 2b). All crosslinked intermolecular lysines are present in the alpha helical region (Fig. 2a). We next removed the flexible loops and fused the shortened CTD (residues 258-301) to SUMO, a monomeric protein with an estimated MW of 17.8 kDa on SEC (Fig. 2c, 2d). We observed that, in comparison with the monomeric SUMO protein, the shortened CTD of AAGAB was sufficient to drive the tetramerization of the fusion protein (~ 56.4 kDa on SEC) (Fig. 2d), confirming that CTD is a *bona fide* tetramerization domain. We therefore referred to the CTD as a tetramerization domain (TD).

### Crystal Structure of AAGAB TD reveals a dimer of dimer assembly

We next crystallized AAGAB TD and solved its atomic structure using single-wavelength anomalous dispersion of the selenomethionyl protein (Se-SAD) (Fig. 3a). The crystal structure was refined to 2.1 Å with R_work_/R_free_ = 21.2/25.9% (Supplementary Table 2). The Se-Met sites enabled us to register all residues without ambiguity. The high-quality main-chain and side-chain electron density allowed us to confirm the assignment with high confidence (Supplementary Fig. 1).

**Fig. 3.**
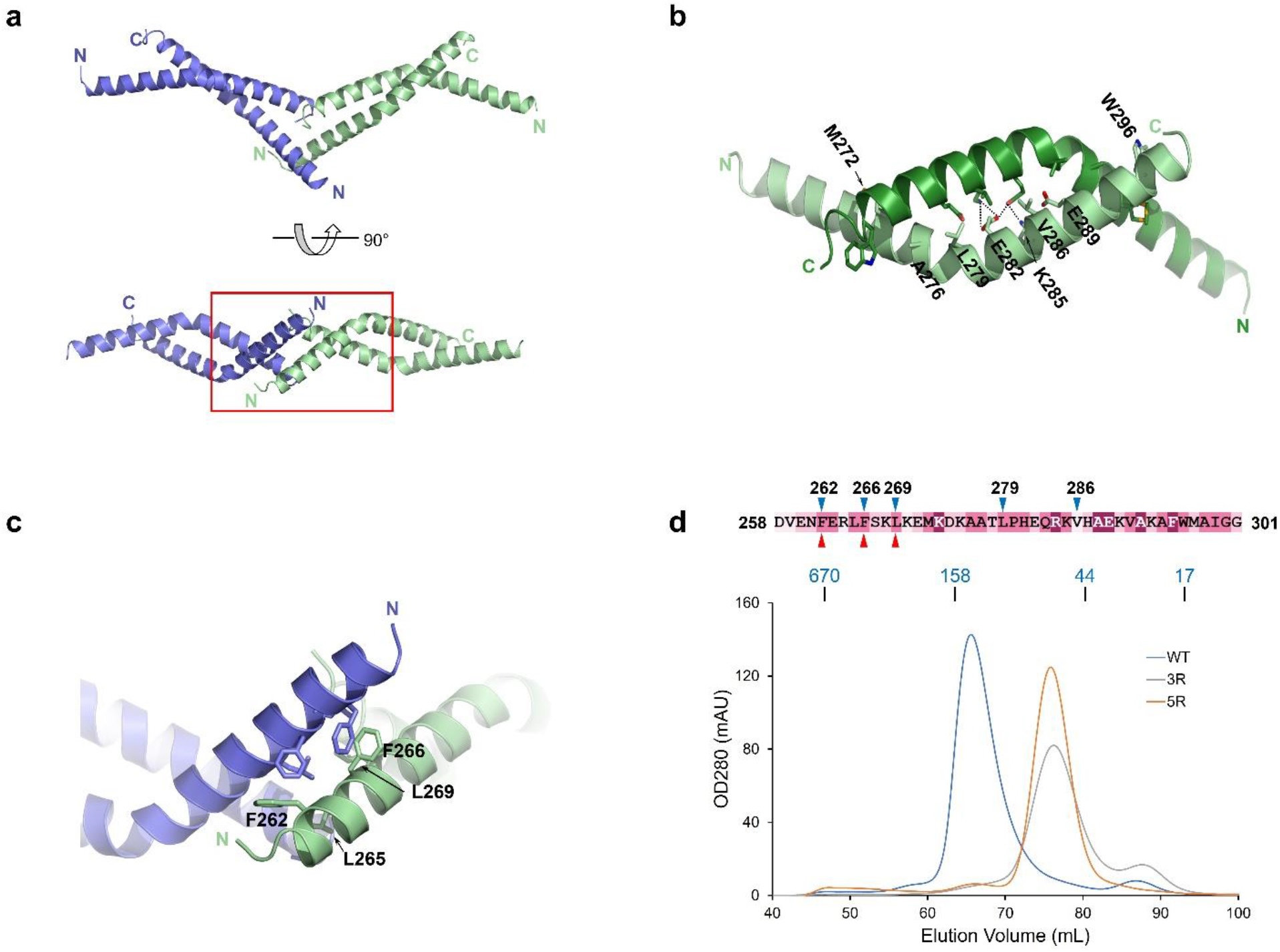
Crystal structure of AAGAB TD reveals a dimer of dimer. **a** Cartoon representation of the AAGAB TD tetramer in two orthogonal views. One dimer is colored in blue while the other is colored in pale green. The N- and C-termini are marked for one molecule in each dimer. **b** The interaction details of the dimer interfaces. Since the interfaces are symmetrical, only one set of residues involved in dimerization are labeled for clarity. **c** The tetramer interface in the zoom-in view of the red box in **a**. **d** SEC profiles of the AAGAB-WT, −3R and −5R mutants from a HiLoad 16/600 Superdex 200 column. Elution positions of protein standards with known MW are marked on the top. The mutated residues of AAGAB-3R and −5R are marked by red and blue arrowheads on the TD sequence, respectively.

The crystal structure of AAGAB TD revealed a tetrameric assembly, fully consistent with our biochemical data. The TD of AAGAB folds into a continuous alpha helix with a bend introduced by P280, adopting a boomerang-like configuration (Fig. 3a). Two TDs run antiparallelly to form a homodimer (Fig. 3b). The two concave sides of the boomerangs interlock each other through an extensive dimer interface, burying a total surface area of 1777.2 Å^2^. The dimer interface is mainly mediated by hydrophobic residues and hydrophobic moieties of charged residues (Fig. 3b). We also observed a few polar interactions in this region. E282 forms a hydrogen bond with another E282 from the other chain in the dimer, while simultaneously forming a salt bridge with K285 also from the other chain in the dimer (Fig. 3b). Two TD dimers further juxtapose in a head-to-head fashion to form a TD tetramer. The tetramerization interface, with a combined buried surface area of 653.9 Å^2^, is formed by the outer side of the N-terminal region of TD. F262 and F266 on the two molecules constitute the core of the predominantly hydrophobic tetramer interface, while L265 and L269 reinforce the interface from periphery (Fig. 3c). These structural analyses indicate that the AAGAB tetramer is formed by dimerization of two dimers (Fig. 3a, 3c).

To test the functional role of the tetramerization interface observed in the crystal structure, we introduced triple mutations (F262R/F266R/L269R) into the tetramer interface of the FL AAGAB protein, designated AAGAB-3R. Indeed, the peak of AAGAB-3R in the SEC profile shifted from the tetramer position to a dimer position (Fig. 3d), indicating a complete disruption of tetrameric assembly. We also introduced two additional point mutations (L279R/V286R) into the dimer interface of the AAGAB-3R protein, aiming to further disrupt the dimer into monomers. However, the peak of this quintuple mutant protein (designated AAGAB-5R) in the SEC profile was similar to that of the dimeric AAGAB-3R mutant (Fig. 3d), suggesting that the two point mutations at the dimer interface were insufficient to disrupt the dimer. These data are consistent with a much larger buried interface area on the dimer interface than the tetramer interface revealed in the structure. We chose not to introduce additional point mutations to avoid disruption of protein folding. Instead, we focused on the 3R and 5R mutants in the subsequent studies.

Overall, the tetrameric assembly uncovered in the structure agrees well with the biochemical data of FL AAGAB (WT and mutants) and the TD domain in solution.

### AAGAB interacts with AP1γ subunit and AP2α subunit with a 1:1 molar ratio

To regulate the assembly of AP1 and AP2 adaptors, AAGAB first interacts with the γ subunit of AP1 or the α subunit of AP2 to form AAGAB:γ or AAGAB:α binary complexes (Wan *et al*., 2021). In these binary complexes, AAGAB stabilizes the γ or α subunit and prepares them for subsequent assembly with other AP1/2 subunits (Gulbranson *et al*., 2019). Next, we expressed and purified recombinant AAGAB:γ and AAGAB:α binary complexes from *E. coli*. The SEC peak elution position of the tag-free AAGAB:γ (trunk domain, residues 1-595) binary complex yielded a calculated molecular weight of 149 kDa (Fig. 4a), suggesting a 1:1 molar ratio of the AAGAB:γ binary complex. The band intensities of γ and AAGAB from the peak fractions are also consistent with a molar ratio of 1:1 (Fig. 4b). We therefore conclude that the AAGAB:γ binary complex contains one copy of each protein.

**Fig. 4.**
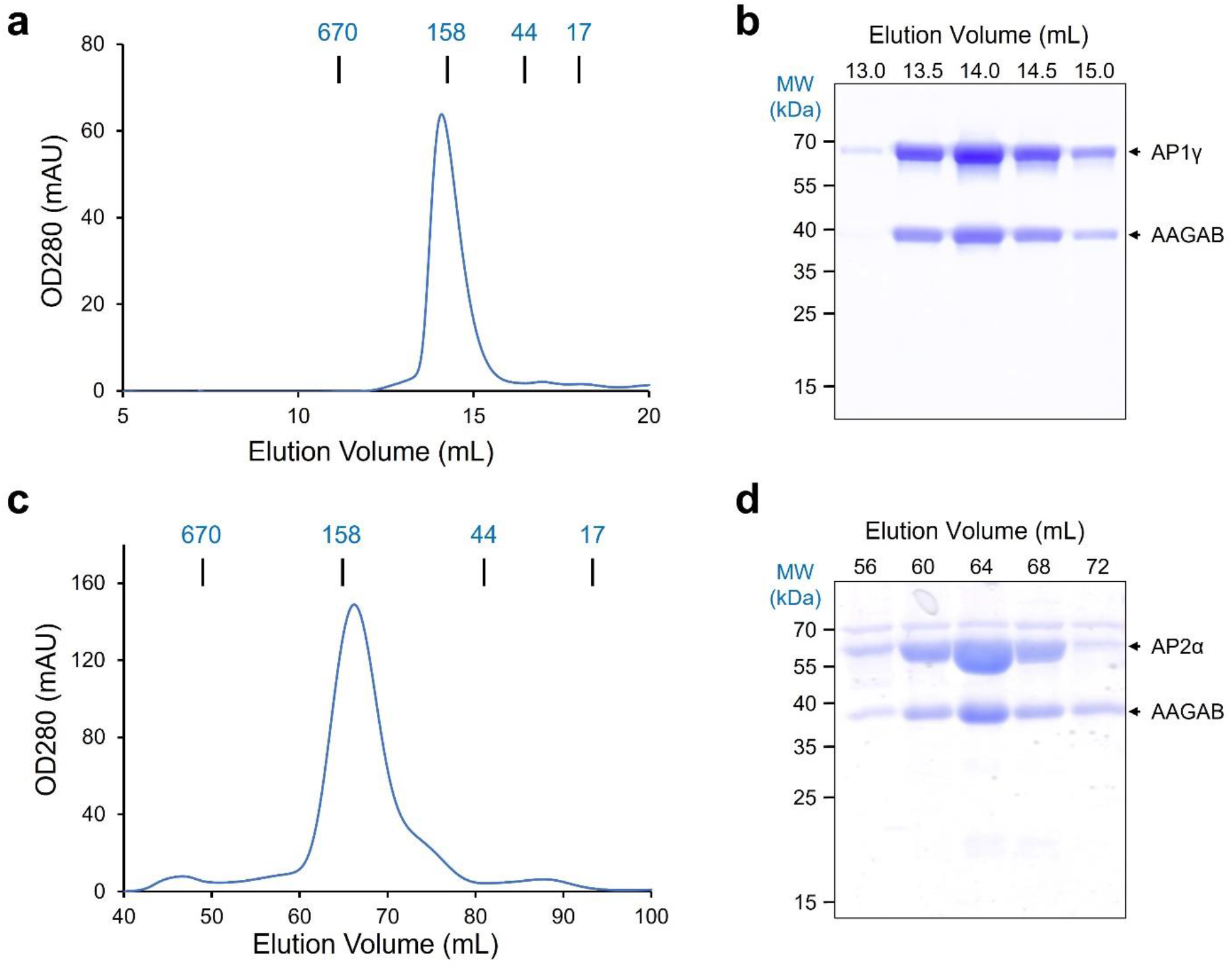
AAGAB interacts with AP1γ and AP2α subunits with a 1:1 molar ratio. **a** SEC profile of the AAGAB:AP1γ (trunk domain, residues 1-595) binary complex from a Superdex 200 Increase 10/300 column. Elution positions of protein standards with known MW are marked on the top. **b** AAGAB and AP1γ proteins in peak fractions of **a** were analyzed by SDS-PAGE and stained by Coomassie blue. **c** SEC profile of AAGAB:AP2α (trunk domain, residues 1-621) binary complexes from a HiLoad 16/600 Superdex200 column. Elution positions of protein standards with known MW are marked on the top. **d** AAGAB and AP2α proteins in peak fractions of **c** were analyzed by SDS-PAGE and stained by Coomassie blue.

When co-expressed in *E. coli,* His_6_-SUMO-AAGAB interacted with GST-tagged α subunit (trunk domain, residues 1-621) in a GST pull-down assay (Gulbranson *et al*., 2019). To further characterize the AAGAB:α binary complex, we removed both the His_6_-SUMO and GST tags after purification using tandem Ni-NTA beads and GSTrap affinity chromatography. The purified tag-free AAGAB:α binary complex exhibited excellent homogeneity and purity on SEC and Coomassie blue-stained SDS-PAGE gel (Fig. 4c, 4d), confirming that AAGAB formed a stoichiometrically stable complex with the α subunit in solution. Interestingly, the SEC profile of the AAGAB:α binary complex exhibited a molecular weight of 139 kDa, suggesting a 1:1 molar ratio. The Coomassie blue-stained SDS-PAGE gel of AAGAB:α peak fractions also suggests that the AAGAB:α binary complex adopts a 1:1 stoichiometry.

Taken together, AAGAB forms a stoichiometric binary complex with the γ subunit of AP1 or the α subunit of AP2 with a 1:1 ratio. These data indicate that AAGAB undergoes a tetramer-to-monomer transition when it engages in AP1 and AP2 binding.

### TD mediates the interaction of AAGAB with the γ subunit of AP1 adaptor and the α subunit of AP2 adaptor

We next sought to pinpoint the region(s) in AAGAB that interacts with AP1γ subunit and AP2α subunit. We co-expressed GST-tagged γ subunit (trunk domain, residues 1-595) with full-length or truncated AAGAB bearing an N-terminal His_6_-SUMO tag in *E. coli*. Consistent with SEC results (Fig. 4a, 4b), co-expression with FL AAGAB strongly increased soluble GST-tagged γ subunit through the formation of the AAGAB:γ binary complex (Fig. 5a). When GD was co-expressed, however, GST-tagged γ subunit was not stabilized even when GD expressed abundantly (Fig. 5a). In contrast, soluble AP1γ subunit was markedly enhanced when co-expressed with AAGAB TD (residues 228-315) and was able to pull down AAGAB TD. These results suggest that AAGAB TD, but not GD, binds and stabilizes the AP1γ subunit. We also tested a shortened version of AAGAB TD (residues 258-301). Again, soluble GST-tagged γ was strongly enhanced through a direct interaction with AAGAB TD (Fig. 5a). Thus, we conclude that AAGAB TD directly recognizes and stabilizes the γ subunit of AP1 adaptor.

**Fig. 5.**
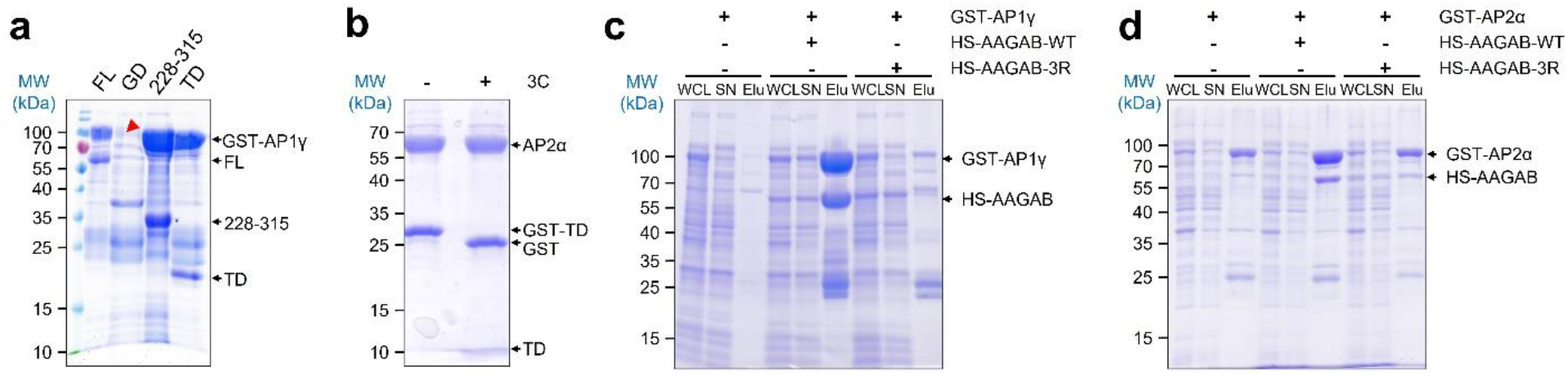
AAGAB TD directly recognizes and stabilizes AP1γ and AP2α subunits. **a** Coomassie blue-stained gel showing the binding of GST-AP1γ (trunk domain, residues 1-595) to His_6_-SUMO tagged FL AAGAB and individual domains. Tetramerization domain (TD): residues 258-301 of AAGAB. Note that there is no specific soluble protein in lane 2 (indicated with a red arrowhead). **b** Coomassie blue-stained gel showing the binding of AP2α subunit (trunk domain, residues 1-621) to GST-AAGAB TD. The identity of GST-AAGAB TD is confirmed by 3C protease treatment to remove the GST tag. **c**-**d** Coomassie blue-stained gels showing the soluble GST tagged AP1γ subunit (**c**) and AP2α subunit (**d**) when they are expressed alone or co-expressed with His_6_-SUMO (HS) tagged AAGAB-WT or 3R mutant. WCL, whole cell lysate. SN, supernatant. Elu, elution.

To test whether TD also binds the α subunit of AP2 adaptor, we co-expressed GST-tagged TD and His_6_-SUMO tagged α subunit (trunk domain, residues 1-621) in *E. coli*. After tandem Ni- and GST-affinity chromatography and Ulp1 protease treatment to remove His_6_-SUMO from the α subunit, we were able to pull down AP2α subunit bound to GST-AAGAB-TD (Fig. 5b). Thus, AAGAB TD directly recognizes and stabilizes both the γ and α subunits.

Next, we sought to define the binding interface of the AAGAB-AP1/2 binary complexes. As expected, co-expression with AAGAB-WT substantially increased the levels of soluble AP1γ (Fig. 5c). Interestingly, the levels of soluble AP1γ were not enhanced when it was co-expressed with the AAGAB-3R mutant, which was defective in tetramerization, and an AAGAB:γ binary complex was not observed (Fig. 5c). Since expression levels of AAGAB-WT and AAGAB-3R were comparable (Supplementary Fig. 2), we conclude that the triple mutations F262R/F266R/L269R, which disrupt AAGAB tetramerization, also abolish the interaction between AAGAB and AP1γ subunit. Similarly, AAGAB-WT, but not the AAGAB-3R mutant, bound and stabilized AP2α subunit when the proteins were co-expressed (Fig. 5d). Taken together, these data demonstrate that AAGAB TD directly binds and stabilizes the γ and α subunits, and the AAGAB tetramerization interface is also involved in the interactions.

### TD mutations disrupt AP1- and AP2-dependent membrane trafficking in the cell

To examine the physiological relevance of our structural and biochemical findings, we determined how the TD mutations affect AAGAB function in the cell. In *AAGAB* KO cells, AP1 and AP2 adaptors are lost, disrupting both AP1- and AP2-mediated trafficking of membrane proteins (Gulbranson *et al*., 2019; Wan *et al*., 2021). As a result, surface levels of transferrin receptor (TfR) were strongly elevated in *AAGAB* KO HeLa cells and the phenotype was rescued by expression of a WT *AAGAB* gene (Fig. 6a, 6b) (Gulbranson *et al*., 2019; Wan *et al*., 2021). By contrast, we observed that expression of the AAGAB-3R or −5R mutant did not restore TfR surface levels (Fig. 6a, 6b). Likewise, deletion of the TD (AAGABΔTD) also abolished the ability of AAGAB to restore TfR surface levels (Fig. 6a, 6b). We also examined the effects of the AAGAB mutations on stabilization of endogenous AP1 and AP2 subunits. We observed that expression of WT *AAGAB* in *AAGAB* KO cells markedly increased the levels of AP1γ subunit and AP2α subunit whereas none of the AAGAB mutants was able to enhance their expression (Fig. 6c, Supplementary Fig. 3). The AAGAB-3R and −5R mutants increased the levels of AP1σ subunit as well as WT AAGAB (Fig. 6c), implying that TD is not involved in σ binding. While 3R and 5R mutants displayed comparable expression levels as WT AAGAB, AAGABΔTD was expressed at a significantly lower level despite similar mRNA levels (Fig. 6c, Supplementary Fig. 3), suggesting TD is required for AAGAB stability. Consistently, all tested adaptor subunit levels remained low when AAGABΔTD was expressed (Fig. 6c). These results are consistent with our *in vitro* data and demonstrate that TD mediates the binding of AAGAB to AP1γ and AP2α subunits in the cell.

**Fig. 6.**
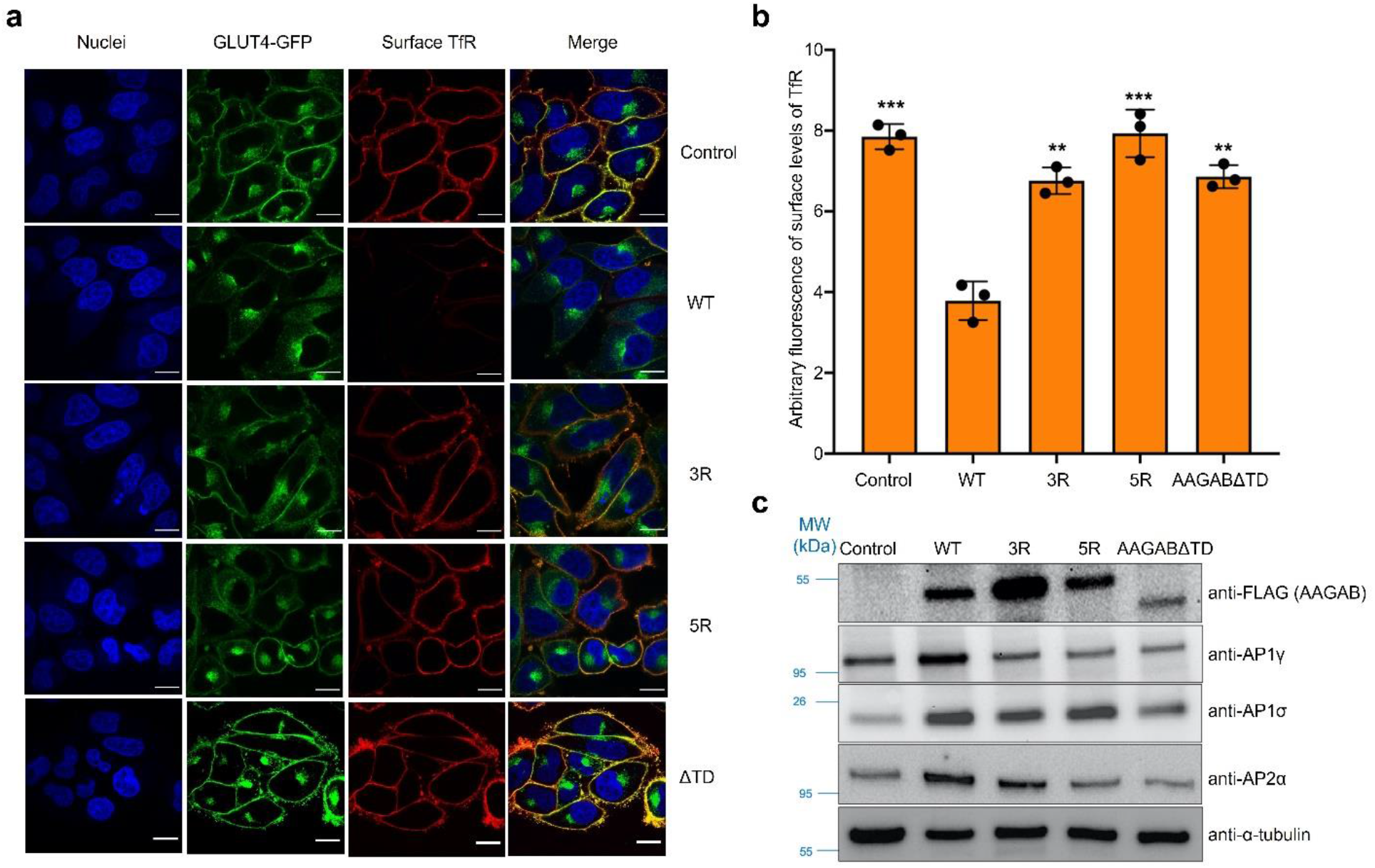
TD mutations disrupt AP1- and AP2-dependent clathrin-mediated trafficking. **a** Representative confocal microscopy images showing surface levels of GLUT4 and TfR in *AAGAB* KO HeLa cells expressing WT or mutant AAGAB. Surface TfR levels of non-permeabilized cells were labeled using anti-TfR antibodies and Alexa Fluor 568-conjugated secondary antibodies (red). Cellular GLUT4 levels were visualized by GFP fluorescence (green). Nuclei were stained with Hoechst 33342 (blue). Images were captured using a 100× oil immersion objective on a Nikon A1 Laser Scanning confocal microscope. Scale bars: 10 μm. **b** Normalized surface levels of TfR in *AAGAB* KO HeLa cells expressing WT or mutant *AAGAB* genes. Surface TfR levels were measured by flow cytometry. Cells were disassociated by Accutase and stained with monoclonal anti-TfR antibodies and APC-conjugated secondary antibodies. APC fluorescence measurements of ~5,000 cells were collected on a CyAn ADP analyzer. Mean APC fluorescence of mutant cells was normalized to that of WT cells. Data are presented as mean ± SD, n=3. \*\**P*<0.01, \*\*\**P*<0.001. P values were calculated using one-way ANOVA. **c** Representative immunoblots showing the expression of the indicated proteins in *AAGAB* KO HeLa cell expressing WT or mutant AAGAB. Control: cells transfected with an empty vector.

## Discussion

Individual subunits and assembly intermediates of a protein complex are often prone to misfolding, aggregation, and degradation. Substrate-specific assembly chaperones like AAGAB stabilize these structures and maintain them at states competent for assembly. In this work, we discovered that in its resting state AAGAB exists as a homo-tetramer. AAGAB tetramerization is mediated by a conserved C-terminal TD. Intriguingly, the same TD interacts with AP1γ and AP2α subunits in a 1:1 molar ratio and is required for stabilizing these subunits both *in vitro* and *in vivo*. Moreover, the same hydrophobic residues (F262, F266, and L269) are involved in both AAGAB tetramerization and interactions with AP1γ or AP2α. We propose a model that AAGAB exists in the tetrameric state prior to AP1/2 binding, and tetramerization shields the hydrophobic AP1/2-binding interface from the aqueous environment and thus stabilizes the substrate-free AAGAB proteins. To engage in AP1/2 binding, an AAGAB tetramer dissociates into monomers and each of them binds one AP1γ or AP2α subunit using the same TD, leading to formation of binary complexes. This model nicely explains how AAGAB itself is stabilized prior to AP1/2 binding.

How is the tetramer-to-monomer transition of AAGAB achieved? AAGAB may exist in a tetramer-monomer equilibrium, with monomeric AAGAB being the binding-competent form. As such, presence of AP1/2 subunits spontaneously drives dissociation of AAGAB tetramers. However, given the highly hydrophobic nature of the interfacial residues, monomeric or dimeric AAGAB may not be stable, which is consistent with the fact that we did not observe substantial dissociation of AAGAB in serial dilution experiment (Fig. 1a). Thus, we postulate that dissociation of AAGAB tetramers into monomers is triggered by and/or coupled to binding to AP1γ/AP2α (Fig. 7a). It is also possible that unidentified cellular factors exist to facilitate AAGAB dissociation into monomers.

**Fig. 7.**
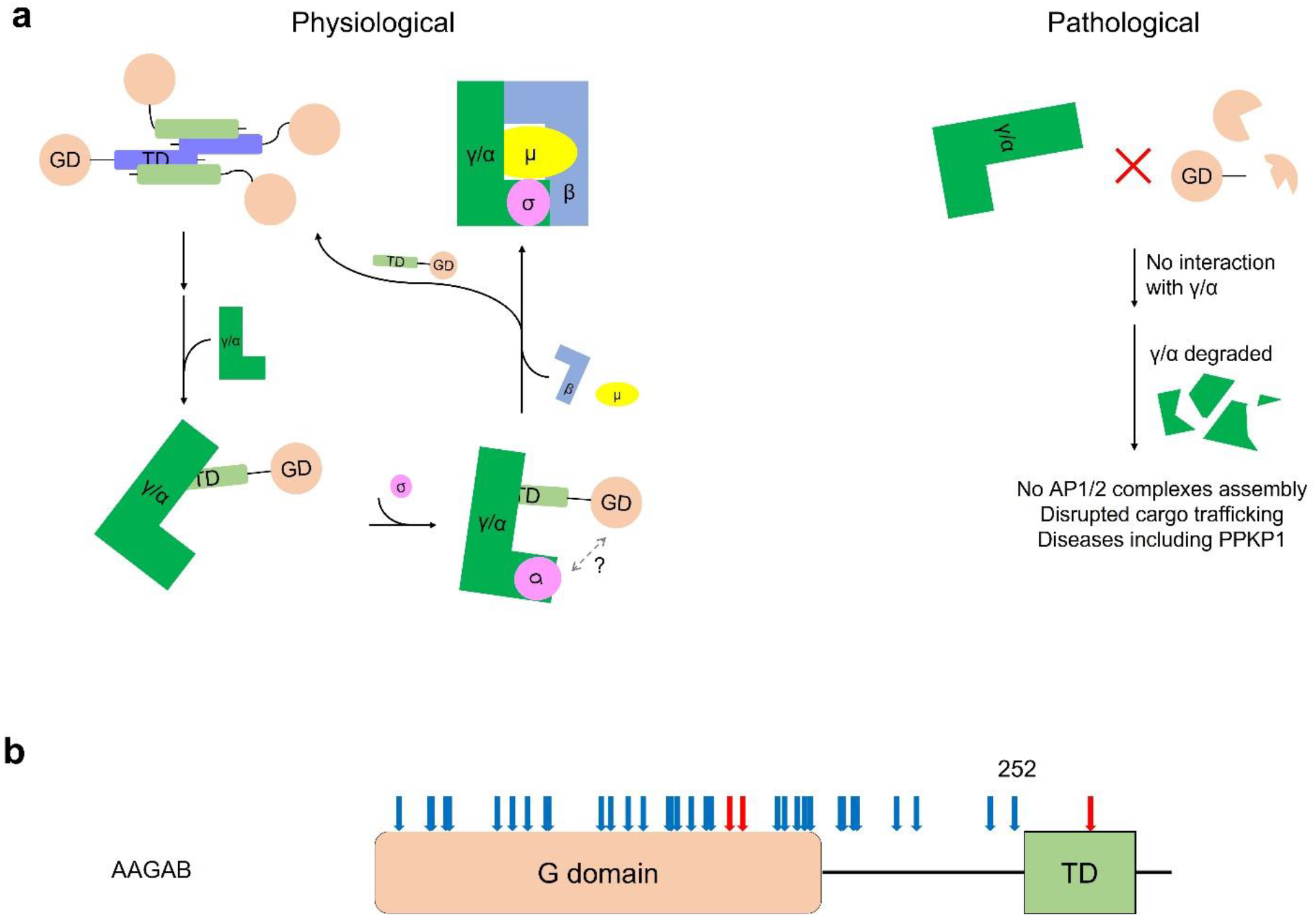
Model of AAGAB-assisted AP1/2 adaptor assembly. **a** AAGAB itself exists as a tetramer in its resting state. An AAGAB tetramer dissociates into monomers upon binding to AP1γ or AP2α subunit using its TD domain. In the AAGAB:γ and AAGAB:α binary complexes, AAGAB stabilizes the AP1/2 subunits and prepare them for subsequent association with other subunits ; σ subunit joins in to form an AAGAB:γ/α:σ ternary hemicomplex; β and μ subunits then displace AAGAB to form the fully assembled AP complexes, while AAGAB returns to the tetramer reservoir. Under pathological conditions, AAGAB lacking the TD domain is unable to interact with and stabilize AP1γ or AP2α subunit, leading to degradation of AP1γ or AP2α subunit, no assembly of AP complexes, and disrupted membrane trafficking. **b** Reported PPKP1-causing mutations in the *AAGAB* coding region are mapped onto AAGAB domain diagram. Nonsense and frameshift mutations are depicted by blue arrows whereas missense mutations are marked by red arrows.

Among a total of 36 known distinct disease mutations reported in the *AAGAB* coding region, 33 are nonsense or frameshift mutations that result in deletion of the entire TD of AAGAB (Fig. 7b) (Giehl *et al*., 2012; Hasegawa et al., 2021; Pohler *et al*., 2012). While these point mutations may also impact AAGAB activity through other mechanisms such as nonsense-mediated mRNA decay, our data clearly demonstrate that these *AAGAB* mutants lose the ability to bind AP1γ and AP2α subunits (Fig. 7a). Heterozygous mutations of *AAGAB* in PPKP1 patients reduce functional AAGAB proteins by half, and the consequences on clathrin-mediated trafficking appear to be restricted to certain tissues such as the skin. Alteration of membrane protein homeostasis in the skin is likely one of the causes of PPKP1. Notably, mutations in genes encoding AP1 subunits such as *AP1B1, AP1S1*, and *AP1S3* are reported to underlie skin disorders (Alsaif et al., 2019; Boyden et al., 2019; Montpetit et al., 2008; Setta-Kaffetzi et al., 2014), indicating malfunction of AP1-mediated cargo trafficking may be the main cause of PPKP1. Intriguingly, one of the three *AAGAB* missense mutations causing PPKP1 is within TD (E282K), and the residue E282 lies at the dimer interface of TD (Fig. 3b).

The bound AAGAB assists in recruitment of small σ subunits and facilitate AP1γ:σ and AP2α:σ interactions (Gulbranson *et al*., 2019; Wan *et al*., 2021). Ultimately, AAGAB is replaced by β and μ subunits, resulting in functional heterotetrameric AP1 and AP2 adaptor complexes and return of AAGAB to the reservoir (Fig. 7a). Although we have reached in-depth understanding of the function for AAGAB C-terminal TD region, the exact function of its conserved N-terminal GD region remains unknown. The crystal structure of GD from the putative yeast homologue Irc6p was solved and found to adopt a genuine GTPase fold (Gorynia et al., 2012). However, Irc6p lacks GTP binding motifs and barely binds to GTP (Gorynia *et al*., 2012). In human AAGAB GD, the GTP binding motifs are also disrupted, indicating it may not possess any GTP binding or hydrolysis capacity either. Further biochemical and structural studies are needed to uncover the function of GD and reveal the molecular basis of AAGAB:σ interactions.

A future research direction is to establish the structural basis of the interactions between AAGAB and AP1/2 complex subunits. This information will likely reveal certain differences between AAGAB-AP1 and AAGAB-AP2 binding interfaces and explain why AAGAB does not regulate the AP3 adaptor complex (Wan *et al*., 2021). Additionally, such structural information could potentially guide us to selectively disrupt AAGAB interaction with one AP complex without affecting the other by introducing well positioned mutations. In this way, we will be able to dissect the roles of AP1 and AP2 adaptor complexes in PPKP1 pathogenesis. Importantly, should our pathogenesis model prove valid, chemical chaperones that mimic TD function and stabilize γ or α subunit could potentially rescue their expression, restore clathrin-mediated membrane trafficking, and reinstate membrane protein homeostasis. Such chemical chaperones may represent new therapeutics to treat PPKP1 patients.

## Methods

### Construction of expression plasmids

DNA sequences encoding full-length human AAGAB (residues 1-315) and individual domains (residues 1-177, 252-315, 258-301) were generated using a standard PCR-based cloning strategy. They were all inserted between *BamH*I and *Sal*I sites at the first multiple cloning site in a modified pRSFDuet-1 vector with a His_6_-SUMO (small ubiquitin-related modifier) tag at the N-terminus. N-terminally His_6_-SUMO tagged mouse AP2α (residues 1-621, which shares a 99% sequence identity with human AP2α) was generated in a similar method. AAGAB mutants were generated by a two-step PCR-based overlap extension method.

For GST tagged recombinant protein expression, FL AAGAB (residues 1-315), AAGAB TD (residues 258-301), and AP2α (residues 1-621) genes were subcloned into the pGEX-6P-1 vector, whereas AP1γ (residues 1-595) was subcloned into the pGEX-4T-3 vector.

For transient expression in mammalian cells, the human *AAGAB* gene was subcloned into the p3xFLAG 7.1 vector, yielding the p3xFLAG7.1-AAGAB-WT plasmid. p3xFLAG7.1-AAGAB-3R and p3xFLAG7.1-AAGAB-5R mutants were generated using bacterial expression plasmids as PCR templates and subsequently subcloned into the p3xFLAG7.1 vector. p3xFLAG7.1-AAGABΔCTD was made by inserting a stop codon after residue 258 in the plasmid p3xFLAG7.1-AAGAB-WT.

### Protein sequence analysis

The sequence analysis of protein FL AAGAB was performed by the online server (https://predictprotein.org) (Bernhofer et al., 2021). It not only generated the secondary structure prediction for FL AAGAB, but also reported the conserved regions in AAGAB using the ConSurf server (https://consurf.tau.ac.il) (Berezin et al., 2004).

### Expression and purification of recombinant proteins

All recombinant proteins were expressed in BL21(DE3) *E. coli* cells in LB medium supplemented with proper antibiotics. The cells were grown at 37 °C and induced by 0.4 mM IPTG when OD600 reached a value of 0.6-0.8, followed by culturing overnight at 20 °C. Cells were harvested by centrifugation at 4,000 × g for 20 min. Cell pellets were lysed by sonication followed by centrifugation at 15,000 × g at 4 °C for 60 min to remove cell debris. Proteins were purified from the soluble fractions by nickel affinity chromatography, followed by SUMO protease Ulp1 treatment (w:w 1:1000) overnight at 4 °C to cleave the His_6_-SUMO tag. The His_6_-SUMO tag was later removed from samples via a second round of nickel affinity chromatography. Proteins were further purified by size exclusion chromatography using a HiLoad 16/600 Superdex-200 prep grade (PG) column (Cytiva #28-9893-35) in the HBS buffer (50 mM HEPES buffer pH 7.5, 150 mM NaCl, and 2 mM β-mercaptoethanol). Peak fractions containing the desired proteins were assessed by SDS-PAGE for purity, pooled and concentrated using Amicon Ultra Centrifugal filters (MilliporeSigma) with a 10 kDa or 50 kDa MW cut-off. Concentrated proteins were aliquoted, flash frozen in liquid nitrogen, and stored at −80 °C.

The L-SeMet derivatized protein was produced using the feedback-inhibition of methionine synthesis pathway. L-SeMet was added to bacterial cell cultures at IPTG induction to a final concentration of 0.1 mg/mL. SeMet-containing AAGAB TD was purified in the same way as the native protein. β-mercaptoethanol (5 mM) was maintained throughout protein purification to prevent oxidation of SeMet.

For AAGAB:AP2α binary protein complex purification, GST-AAGAB (Ampicillin resistant) and His_6_-SUMO-AP2α (Kanamycin resistant) plasmids were co-transformed into BL21(DE3) *E. coli* cells to express the complex. The complex was expressed in the same way as individual proteins. After cell lysis and centrifugation, the binary complex was first purified by nickel affinity chromatography, followed by overnight Ulp1 treatment (w:w 1:1000) at 4 °C to cleave the His_6_-SUMO tag. The Ulp1 treated protein sample was loaded directly onto a prepacked GSTrap column (GE Healthcare). Protease 3C was added to the eluted GST-AAGAB:AP2α binary complex to cleave the GST tag, which was removed by a second GSTrap column. The tag-free binary protein complex was finally purified by size-exclusion chromatography using a HiLoad 16/600 Superdex-200 PG column (Cytiva #28-9893-35) in the HBS buffer. AAGAB TD:AP2α binary complex was expressed and purified using a similar method.

### Crystallization, data collection, structure determination, and refinement

AAGAB TD crystals were grown using the hanging-drop vapor-diffusion method by mixing the protein (27 mg/mL in HBS buffer) with an equal volume of reservoir solution containing 30% glycerol, 0.5 M ammonium phosphate (Hampton Research) at 16 °C. The shining diamond like crystals started to show up after overnight incubation and reached full size within a week. All crystals were flash frozen in liquid nitrogen without any additional cryo-protectant.

Diffraction datasets were collected on 22-ID and 22-BM beamlines (SER-CAT) at Advance Photon Source (APS), Argonne National Laboratory and AMX (17-ID-1) and FMX (17-ID-2) beamlines at National Synchrotron Light Source II (NSLS-II), Brookhaven National Laboratory.

The datasets of human AAGAB TD were indexed, integrated, and scaled using HKL2000 package (Otwinowski, 1997). The crystal belongs to space group P6_1_22 (a = b = 47.539 Å, c = 191.394 Å, α = β = 90°, γ = 120°) and contains 2 molecules per asymmetric unit. The structure was determined by single-wavelength anomalous diffraction (SAD) using data collected at Se-peak wavelength to a resolution of 2.4 Å. The position of selenium atoms was located by the program AutoSol and the initial model was built by the program AutoBuild, which was later extended to 2.1 Å from a native dataset. Further model improvement was carried out with alternate rounds of refinement using Phenix.refine (Liebschner et al., 2019) and model building via COOT (Emsley et al., 2010). The structure refinement was completed with cycles of individual B-factor refinement along with TLS parameters, leading to a final structure with R_work_ of 21.2% and R_free_ of 25.9%. The final model is of good stereochemical quality and Ramachandran plot of the main-chain angles showed 100% of the residues found in the favored region. The data collection and refinement statistics for this structure is listed in Supplementary Table 2.

### Analytical gel filtration

Recombinant FL AAGAB proteins were analyzed on a Superdex 200 increase 10/300 GL column (Cytiva #28-9909-44) with a flow rate of 0.5 mL/min and an injection volume of 0.5 mL. All other protein samples were analyzed on HiLoad 16/600 Superdex 200 PG column (Cytiva #28-9893-35) with a flow rate of 0.8 mL/min and an injection volume of 4 mL. All experiments were performed in the HBS buffer. The columns were calibrated using the Gel Filtration Standard (Bio-Rad #1511901).

### Lysine-specific crosslinking coupled with nano-Liquid Chromatography Mass Spectrometry (nLC-MS)

The purified AAGAB protein was cross-linked with a solution of 1:1 BS3-*d*0: BS3-*d*4 (ThermoFisher #21590 and #21595) crosslinkers. Crosslinking reaction product was separated by SDS-PAGE. The bands corresponding to dimer and tetramer were cut. Cut gel bands were destained, reduced with dithiothreitol (DTT), and digested with trypsin at 37°C overnight. Tryptic peptides were separated by an Easy Nano LC II system (Thermo Scientific). Mobile phases were water with 0.1% formic acid (A) and acetonitrile with 0.1% formic acid (B). A 3-hour gradient (from 5% to 45% B) was performed with a flow rate of 300 nL/min. nLC_eluates were on-line ionized by nano-electrospray ionization (nESI) and detected by a Velos LTQ-Orbitrap Mass Spectrometer (Thermo Scientific). Precursor ions were detected in the Orbitrap with a mass resolution of 60 K, while the data-dependent MS2 of the top 10 most abundant precursor ions were carried out in LTQ. The collected .raw files were converted to .mzXML files for crosslinking data analysis with StavroX, an open-access software (Gotze et al., 2012). Search parameters were used as followed: max 4 trypsin miscleavages, dynamic modification of methionine by oxidation, precursor mass accuracy of better than 5 ppm and fragment ion mass accuracy of better than 0.8 Da. StavroX generated crosslinked peptide list was further manually checked for assignment.

### Pull-down assay using co-expressed recombinant proteins

GST-AP1γ and GST-AP2α were expressed individually or in combination with His_6_-SUMO-AAGAB-WT or −3R plasmids in *E. coli* BL21(DE3) cells. Cells were cultured in 500 mL LB medium with corresponding antibiotics. After harvesting, cells were lysed and centrifuged following the same protocol as described above. The cleared cell lysates were equally split for either GST or nickel affinity pull-down. For GST affinity pull-down, the cleared cell lysate was loaded onto a gravity column with 1 mL bed volume of Glutathione agarose resin (Gold Biotechnology, #G-250-5). The resin was extensively washed, and bound proteins were eluted with a GST elution buffer (50 mM Tris-HCl pH 8.0, 150 mM NaCl, and 30 mM L-Glutathione reduced) in 1 mL fractions. For nickel affinity pull-down, the cleared cell lysate was loaded onto a gravity column with 0.5 mL bed volume of Ni-NTA resin (Qiagen, #30210). The resin was extensively washed, and bound proteins were eluted with a Ni-NTA elution buffer (50 mM Tris-HCl pH 8.0, 150 mM NaCl, and 300 mM imidazole) in 0.5 mL fractions. All elution fractions were analyzed by SDS-PAGE.

### Flow cytometry

HeLa cells were maintained in Dulbecco’s Modified Eagle Medium (DMEM) supplemented with 10% FB essence (FBE; Seradigm, #3100-500) and penicill-instreptomycin (Millipore-Sigma, #P4333). To stain surface TfR, HeLa cells were washed three times with the KRH buffer (121 mM NaCl, 4.9 mM KCl, 1.2 mM MgSO_4_, 0.33 mM CaCl_2_, and 12 mM HEPES, pH 7.0). Cells were then chilled on ice and stained with monoclonal anti-TfR antibodies (DSHB, #G1/221/12) at a final concentration of 0.1 μg/mL and APC-conjugated secondary antibodies (Thermo Fisher Scientific, #17-4015-82) at a final concentration of 0.8 μg/mL. After dissociation from plates using Accutase (Innovative Cell Technologies, #AT 104), APC fluorescence of the cells was measured on a CyAn ADP analyzer (Beckman Coulter). Mean APC fluorescence of mutant cells was normalized to that of WT cells. Data from populations of ~5,000 cells were analyzed using the FlowJo software (FlowJo, LLC, v10) based on experiments run in biological triplicates.

### Immunoblotting

To detect proteins in whole cell lysates, cells grown in 24-well plates were lysed in the SDS protein buffer. Protein samples were resolved on 8% Bis-Tris SDS-PAGE and proteins were detected using primary antibodies and horseradish peroxidase (HRP)-conjugated secondary antibodies. Primary antibody used in this work included monoclonal anti-FLAG antibodies (Millipore-Sigma, #F1804) at a final concentration of 1 μg/mL, polyclonal anti-AP1γ antibodies (Bethyl, #A304-771A) at a final concentration of 1 μg/mL, polyclonal anti-AP2α antibodies (BD Biosciences, #610502) at a final concentration of 1 μg/mL, polyclonal anti-AP1σ antibodies (Bethyl, #A305-396A) at a final concentration of 1 μg/mL and monoclonal anti-α-tubulin antibodies (DSHB, #12G10) at a final concentration of 43 ng/mL.

### Immunostaining and imaging

Cells grown in a 4-compartment 35 mm glass-bottom dish were washed three times with the KRH buffer and fixed using 2% paraformaldehyde. Cells were permeabilized by 0.1%Triton-X100 in PBS. Surface TfR proteins were stained using monoclonal anti-TfR antibodies (DSHB, #G1/221/12) at a final concentration of 0.1 μg/mL and Alexa Fluor 568-conjugated secondary antibodies (Thermo Fisher Scientific, #A11004) at a final concentration of 1 μg/mL. The nuclei were stained with Hoechst 33342 (Thermo Fisher Scientific, #H3570) at a final concentration of 10 μg/mL. Images were captured using a 100× oil immersion objective on a Nikon A1 Laser Scanning confocal microscope and processed using FIJI software (Schindelin et al., 2012).

### Real-time quantitative reverse transcription PCR

Total RNAs were isolated using the RNeasy Mini Kit (Qiagen, #74104), followed by treatment with ezDNAse (Thermo Fisher Scientific, #18091150). First strand complementary DNA synthesis was performed using a SuperScript IV kit (Thermo Fisher Scientific, #18091050). Gene expression was determined by quantitative reverse transcription PCR on Applied Biosystems™7500 Fast-Real Time PCR Detection System using SsoAdvanced Universal SYBR Green Supermix (Bio-Rad, #172-5272) with gene-specific primer sets. The cycle threshold values of a candidate gene were normalized to those of *GAPDH*, a reference gene, and the Δcycle threshold values were calculated. The results were plotted as fold changes relative to the wild type *AAGAB* rescue sample. PCR primers for *AAGAB* and mutants were as follows: 5’-TGACGATGACAAGCTTATGGCT-3’ (forward) and 5’-CGGAAAATACTGAGGAGCAGC-3’ (reverse). PCR primers for *GAPDH* were as follows: 5’-GACAGTCAGCCGCATCTTCT-3’ (forward) and 5’-GCGCCCAATACGACCAAATC-3’ (reverse).

## Supporting information

Supplementary Table 1 AAGAB XL-MS peptides information

Supplementary Table 2 X-ray crystallography data collection and refinement statistics

## Acknowledgments

We thank Drs. James Hurley and Juan Bonifacino for reagents or advice. We thank Micaela Martinez, Marie Chmara, Leanne Diab, and Aditi Krishnan for literature research. We thank Dr. T. “Soma” Somasundaram and beamline scientists at APS 22-ID and 22-BM and NSLSII AMX and FMX (17-ID-1 and 17-ID-2) for assistance in X-ray diffraction data collection. This work was supported by National Institutes of Health grants GM138685 (Q.Y.), AI146330 (Q.Y.), GM126960 (J.S.), DK124431 (J.S.), and AI156560 (S.L.).

## Author Contributions

Y.T., S.L., J.S., and Q.Y. conceived the initial experimental plan. Y.T., R.Y., and B.W. expressed and purified the proteins. B.W. carried out crosslinking reaction and H.H. performed nano-liquid chromatography mass spectrometry. Y.T. and Q.Y. crystallized the TD and determined its structure. Y.T., R.Y. and B.W. carried out co-expression experiments. I.D. and C.W. performed cell-based assays. Y.T. I.D., S.L., J.S., and Q.Y. drafted the manuscript. All authors edited the manuscript.

## Data availability

Coordinates and structural factors for AAGAB TD have been deposited in the Protein Data Bank with the accession code 7TWD.

## Conflict of interest

The authors declare that they have no conflicts of interest with the contents of this article.

## Notes

### Competing Interest Statement

The authors have declared no competing interest.

