## Supplementary Figures 1-3 for "Oligomer-to-Monomer Transition Underlies the Chaperone Function of AAGAB in AP1/AP2 Assembly"

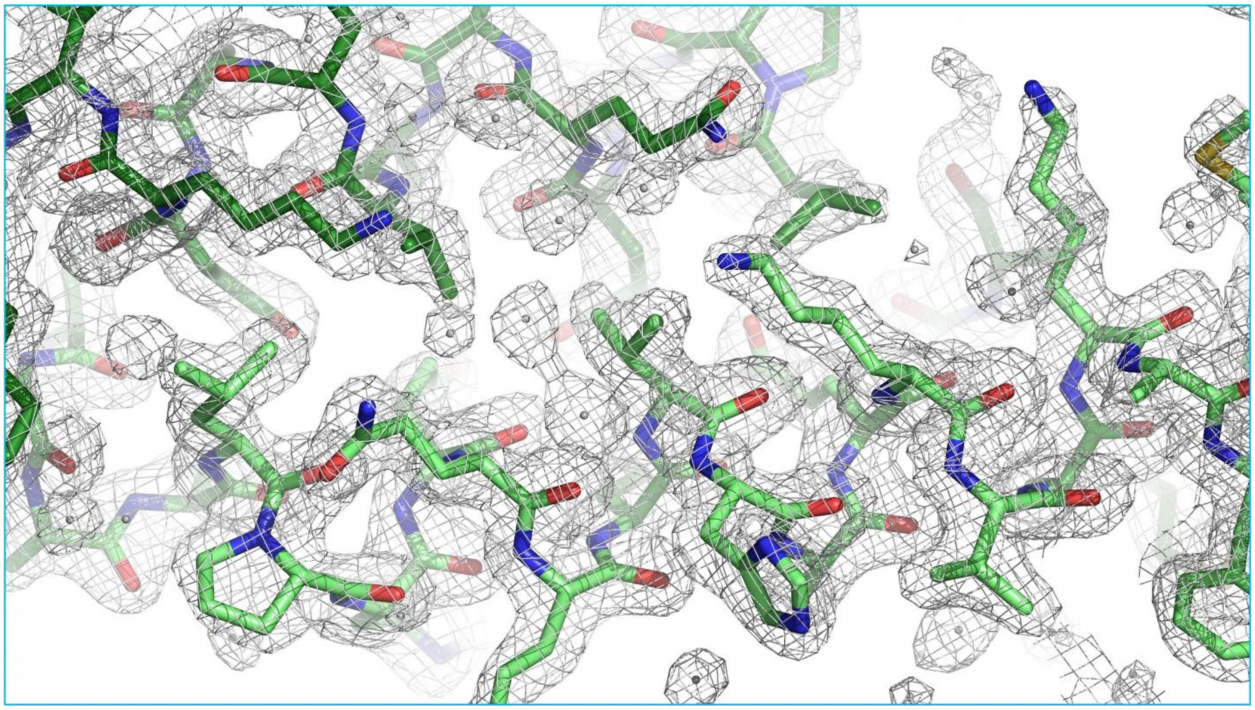

**Suppl. Figure 1** *2Fo-Fc* density map of the TD dimer interface contoured at 1σ level.

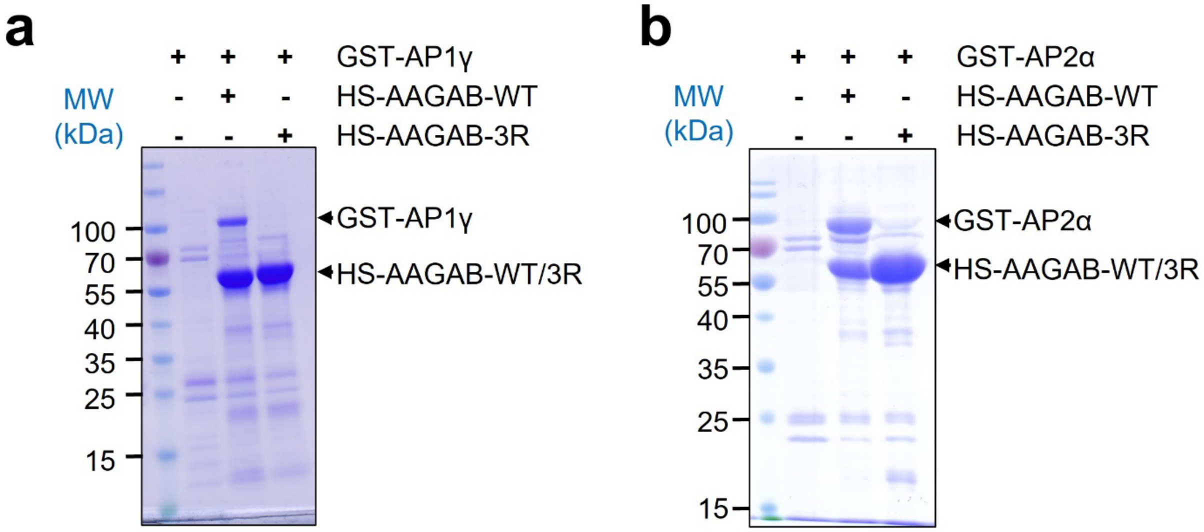


**Suppl. Figure 2** Coomassie blue-stained gels showing the Nickle-NTA eluted soluble proteins when GST tagged AP1γ subunit (**a**) and AP2α subunit (**b**) were expressed alone or co-expressed with $His_6$-SUMO (HS) tagged AAGAB WT or 3R mutant.

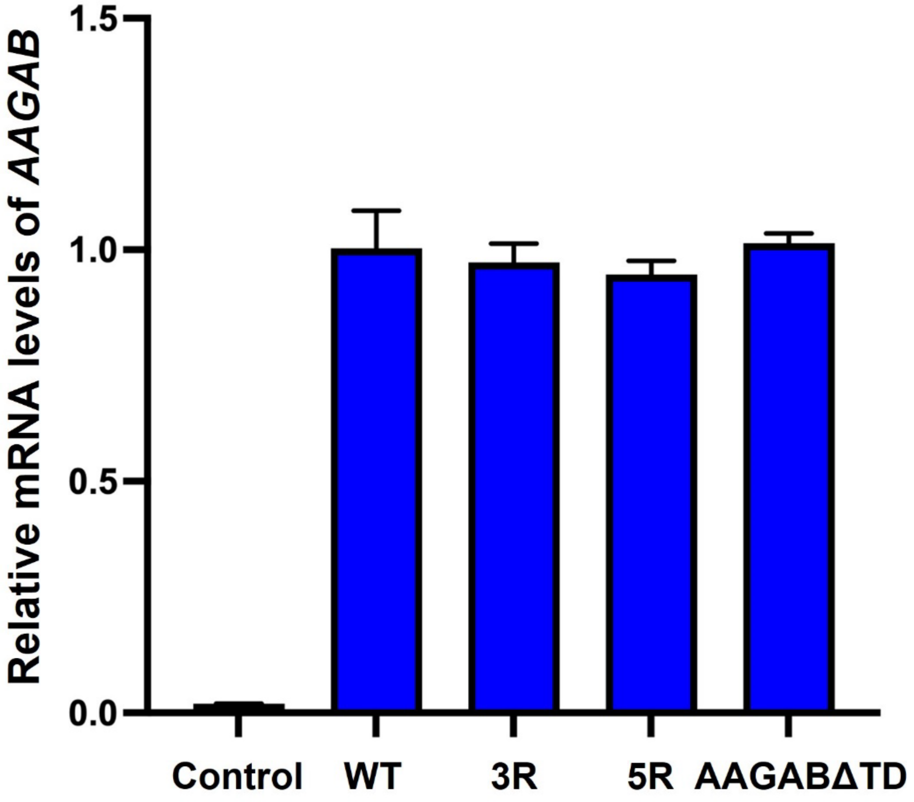


**Suppl. Figure 3** Relative mRNA levels of *AAGAB* were calculated by normalizing the threshold cycles of *AAGAB* to those of *GAPDH*, a gene whose expression remained unchanged in *AAGAB* KO HeLa cells. The fold change was determined by comparing the normalized threshold cycles of mutant *AAGAB*-expressing KO cells to those of WT *AAGAB*-expressing KO cells. Error bars indicate standard deviation (n = 3).
