## Supplementary Table 2 X-ray crystallography data collection and refinement statistics for "Oligomer-to-Monomer Transition Underlies the Chaperone Function of AAGAB in AP1/AP2 Assembly"

**Table 2. Data collection and refinement statistics.**

| | AAGAB TD SeMet | AAGAB TD |
|---|---|---|
| **Data Collection** | | |
| Wavelength (Å) | 0.97929 | 1.000 |
| Resolution range (Å) | 50.00 – 2.40 (2.49 – 2.40) | 50.00 – 2.00 (2.07 – 2.00) |
| Space group | $P6_122$ | $P6_122$ |
| Unit cell (a, b, c, Å) | 47.343, 47.374, 192.670 | 47.539 47.539 191.394 |
| (α, β, γ, °) | 90.0, 90.0, 120.0 | 90.0, 90.0, 120.0 |
| Total reflections | 85083 | 126149 |
| Unique reflections | 5510 (446) | 9351 (900) |
| Multiplicity | 15.4 (6.4) | 13.5 (9.6) |
| Completeness (%) | 97.9 (83.7) | 99.2 (99.7) |
| Mean I/sigma(I) | 19.3 (1.27) | 17.4 (0.93) |
| Wilson B-factor | 39.09 | 22.68 |
| R-merge | 0.143 (0.813) | 0.205 (1.580) |
| R-meas | 0.148 (0.868) | 0.213 (1.658) |
| R-pim | 0.037 (0.287) | 0.057 (0.479) |
| CC1/2 | 0.988 (0.741) | 0.985 (0.380) |
| **Refinement** | | |
| Resolution (Å) | | 37.82 - 2.11 (2.185 - 2.11) |
| Reflections used in refinement | | 7999 (774) |
| Reflections used for R-free | | 802 (78) |
| R-work | | 0.2123 (0.2638) |
| R-free | | 0.2594 (0.3179) |
| Number of non-hydrogen | | 823 |

| | |
|---|---|
| atoms | |
| macromolecules | 739 |
| ligands | 10 |
| solvent | 74 |
| Protein residues | 89 |
| RMS (bonds, Å) | 0.002 |
| RMS (angles, °) | 0.46 |
| Ramachandran favored (%) | 100.00 |
| Ramachandran allowed (%) | 0.00 |
| Ramachandran outliers (%) | 0.00 |
| Rotamer outliers (%) | 0.00 |
| Clashscore | 4.67 |
| Average B-factor | 35.63 |
| macromolecules | 34.08 |
| ligands | 94.50 |
| solvent | 43.18 |
| Number of TLS groups | 1 |

Statistics for the highest-resolution shell are shown in parentheses.
